# Dynamics of membrane tubulation coupled with fission by a two-component module revealed on polymer cushioned bilayer islands

**DOI:** 10.1101/2023.09.17.558094

**Authors:** Soumya Bhattacharyya, Thomas J. Pucadyil

**Affiliations:** Indian Institute of Science Education and Research, Dr. Homi Bhabha Road, Pashan, Pune 411008, Maharashtra, India

**Keywords:** Polymer cushions, BAR domain-containing proteins, dynamin, membrane tubulation, membrane fission, phosphoinositide lipids, lipid-protein interactions

## Abstract

Membrane tubulation coupled with fission (MTCF) is a widespread cellular phenomenon but mechanisms for their coordination remain unclear. This is partly because of the lack of assays to monitor dynamics of membrane tubulation. Using polymer cushioned bilayer islands, we analyze functions of the membrane tubulator Bridging Integrator 1 (BIN1) mixed with the fission catalyst dynamin2 (Dyn2). Our results reveal that this mixture constitutes a minimal two-component module that demonstrates MTCF. MTCF is an emergent property and arises because BIN1 dually functions in facilitating peripheral recruitment while inhibiting membrane binding of Dyn2 in a dose-dependent manner. MTCF is therefore apparent only at high Dyn2 to BIN1 ratios. Because of their mutual involvement in T-tubules biogenesis, mutations in BIN1 and Dyn2 are associated with centronuclear myopathies (CNM) and our analysis links the pathology with aberrant MTCF. Together, our results establish cushioned bilayer islands as a facile template for the quantitative analysis of membrane tubulation and inform of mechanisms that coordinate MTCF.

## Introduction

Live cell imaging has provided valuable insights into the dynamics of membrane tubulation coupled with fission (MTCF) reactions during muscle development, cell migration, vesicular transport and the entry of toxins ^1–7^. A striking example of a regulated tubulation process is apparent during the formation of T-tubules, which are regularly spaced plasma membrane invaginations in cardiac and skeletal muscle cells. Their growth and maturation during muscle development coincides with a change in their protein composition, which allows them to become tightly associated with the sarcoplasmic reticulum and thereby facilitate synchronous excitation-contraction coupling ^8,9^. The muscle-specific isoform (isoform 8) of amphiphysin 2, also called BIN1, and the ubiquitous isoform of dynamin2 (Dyn2) are critical structural components of T-tubules ^10^. BIN1 contains the N-terminal N-BAR domain that bends membranes and the C-terminal Src homology 3 (SH3) domain that interacts with the proline-rich domain (PRD) in dynamins ^11,12^. The clathrin and adaptor protein 2 (AP2) binding motifs are absent in BIN1. Instead, BIN1 contains a polybasic (PI) stretch of amino acids, which confers it with the enhanced ability to bind phosphoinositide lipids ^10,12^. Dynamins are large multimeric GTPases known for their role in membrane fission ^13,14^. Mutations in BIN1 and Dyn2 are linked to autosomal dominant and recessive forms of centronuclear myopathies (CNMs), which are a group of congenital hereditary disorders characterized by disorganized T-tubules leading to uncoordinated force generation and muscle hypotrophy ^15^. The relative levels of BIN1 and Dyn2 are tightly regulated during development, with an increase in BIN1 and a decrease in Dyn2 levels correlating with increased T-tubule densities in muscle cells ^16^. Increased expression of Dyn2 inhibits T-tubule growth during development and overexpression of Dyn2 in myoblasts leads to the fragmentation of BIN1 tubules ^17,18^. Additionally, modulating Dyn2 levels rescues CNM-like defects in disease models ^17,19–22^. Previous reports indicate that BIN1 negatively regulates Dyn2 functions. Thus, BIN1 inhibits dynamin’s stimulated GTPase activity and reduces Dyn2’s ability to vesiculate membranes ^20,23–26^. From a protein design standpoint, if the sole function of BIN1 was to tubulate membranes then just the N-BAR domain and the PI stretch should have sufficed. But BIN1 contains the dynamin-interacting SH3 domain, which implies that fission of BIN1 tubules must at some point be critical for T-tubules biogenesis.

Despite the relevance of membrane tubulation in physiology, a mechanistic understanding of the function of participant proteins has typically relied on end-point electron microscopic (EM) analyses of tubulated liposomes. Such analyses have provided valuable information into the organization of proteins on tubulated membranes. But insights into early intermediates and their dynamics, especially when tubulation is coupled with fission, remain missing. This is because of the lack of suitable membrane templates that are amenable to real-time monitoring of membrane tubulation. Giant unilamellar vesicles (GUVs) are a popular template but their finite reservoir deters membrane tubulation because of heightened membrane tension. Supported bilayers with excess reservoir (SUPER) templates are a work around, but their spherical geometry, as is also the case with GUVs, renders it difficult to track early intermediates during the tubulation reaction ^27^. Planar supported lipid bilayers (SLBs) possess the ideal geometry, but those formed by vesicle fusion on glass are difficult to tubulate because of strong interactions between membrane lipids and the glass surface. This is evident from the dramatic tubulation seen with multilamellar membrane sheets, where the bulk of the membrane reservoir is not in contact with glass ^24,28^. But the use of multilamellar membrane sheets has been restricted to visualizing late stages of membrane tubulation reactions, only after long tubules have been drawn out from the membrane reservoir.

SLBs formed on polymer cushions have reported enhanced diffusion of membrane lipids because the cushion reduces their binding to glass ^29^. Inspired by these findings, we report the utility of polyethylene glycol (PEG)-cushioned planar bilayer islands for the quantitative analysis of membrane tubulation as well as MTCF. Our results with BIN1 and Dyn2 present novel insights into early steps in the tubulation reaction and mechanisms that regulate MTCF. Furthermore, we evaluate functions of CNM-associated mutants in BIN1 and Dyn2 and our results establish a causal link between MTCF and the CNM pathology. Together, our results highlight the potential of cushioned planar bilayer islands for analyzing cellular processes involving membrane tubulation and fission through reconstitution.

## Results

### Cushioned planar bilayer islands as pliable templates for membrane tubulation

Based on previous work from our lab and others, a facile route to forming PEG-cushioned membrane templates is by spreading lipids in an organic solvent as a thin film on a glass coverslip covalently attached to PEG (Fig. S1A). Drying off the solvent and subsequent hydration converts the lipid film into large planar bilayer islands (Fig. S1B) ^30–32^. The vast expanse of the planar bilayer island sometimes displays membrane buds and nanotubes, which appear brighter than the bilayer because of the higher membrane area in these structures (Fig. S1B). Since we were interested in testing BIN1 and Dyn2, we first analyzed their phosphatidylethanolamine ^33,34^. On liposomes containing 5 mol% of different phosphoinositide lipids and 1 mol% of the bifunctional probe in the background of DOPC, BIN1 displays a binding preference in the order of PI(4,5)P_2_ > PI(3,4,5)P_3_ > PI(3,4)P_2_ = PI(3)P > PI(3,5)P_2_>> PI(4P) > PI(5)P (Fig. S1C). The apparent binding affinity of BIN1 for PI(4,5)P_2_ was ∼0.13 μM (Fig. S1D), which falls in the range of previous estimates ^35^. On the other hand, Dyn2 displays a binding preference in the order of PI(4,5)P_2_ > PI(3,4)P_2_ > PI(3,4,5)P_3_ > PI(3,5)P_2_ >> PI(3)P >> PI(4P) = PI(5)P (Fig. S1C). Since PI(4,5)P_2_ is the preferred phosphoinositide lipid for both BIN1 and Dyn2, we resorted to analysis of bilayer islands with 5 mol% of PI(4,5)P_2_ and 15 mol% DOPS in the background of DOPC. The lipid mixture contained 0.5 mol% of the fluorescent lipid analogue Texas Red-DHPE. Flowing 1 μM of the PI(4,5)P_2_ sensor mEGFP-PLCδ PH domain showed uniform binding to the island with no apparent effect on membrane morphology (Fig. S1E). This confirms that PI(4,5)P_2_ is readily accessible and uniformly distributed on the island, and that the templates are stable to protein binding.

Remarkably, flowing 0.2 μM BIN1-mEGFP on to cushioned bilayer islands caused a rapid and profuse tubulation of the island, which in turn caused the bilayer to retract and shrink in size (Fig. 1A and Movie S1). In contrast, flowing a 12-fold higher concentration of 2.5 μM BIN1 on SLBs formed by liposome fusion on glass showed protein binding but no apparent tubulation (Fig. S1F), signifying that cushioned bilayer islands are readily amenable to membrane tubulation. After 10 mins, BIN1 exposed bilayer islands displayed bright tubules that formed an extensive network (Fig. 1B, white arrows), which was also confirmed in scanning electron micrographs (SEM) (Fig. 1C, white arrows). This was because BIN1 tubules showed a tendency to coil around each other (Fig. 1D and Movie S2). Photobleaching the fluorescent lipid probe in a large region of the island displaying a tubular network showed fluorescence recovery in both tubules and the island, indicating that tubules are intact and connected to the island (Fig. S1G, Movie S3).

**Fig. 1.**
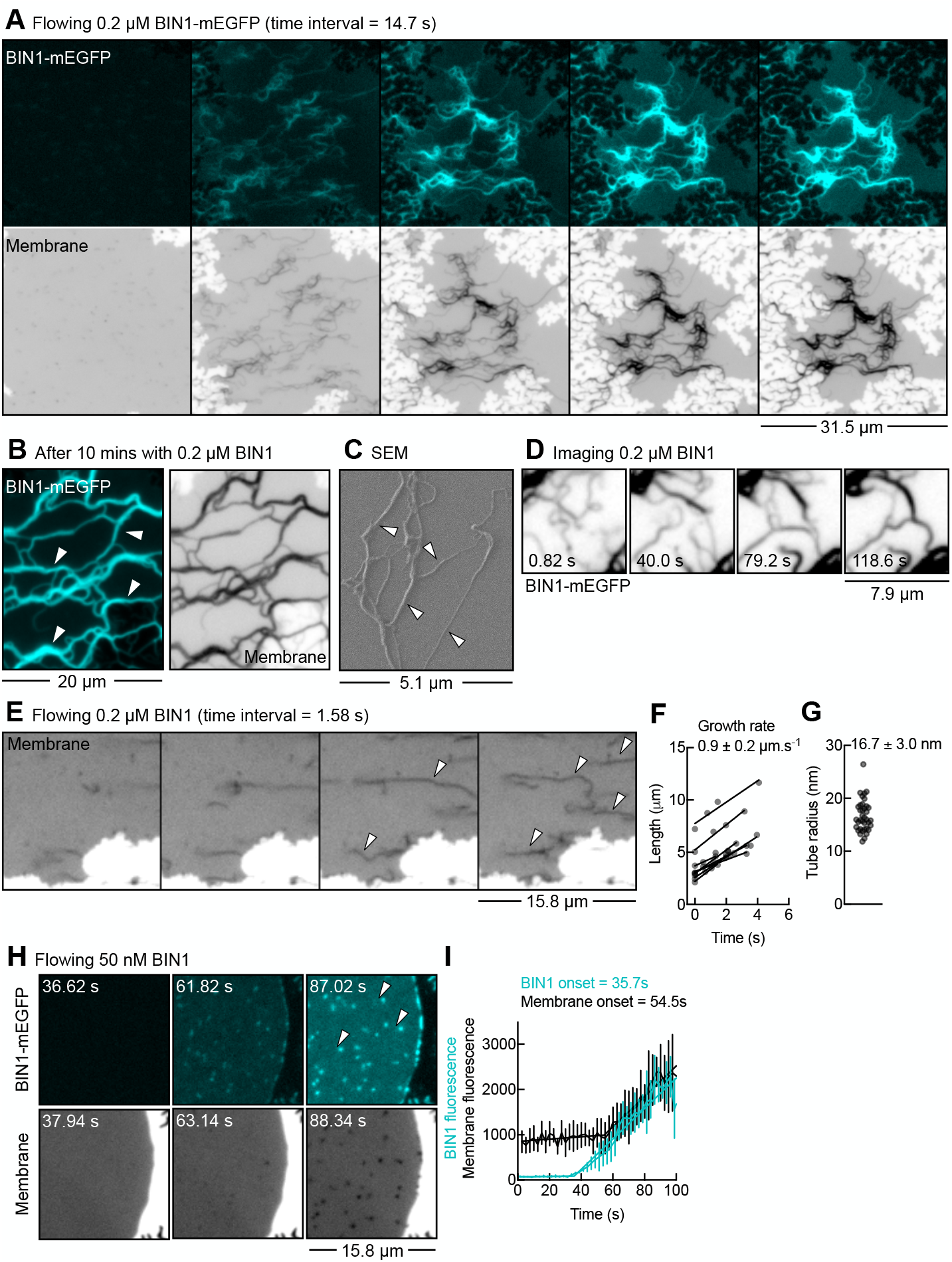
Dynamics of BIN1-induced membrane tubulation. (A) Time-lapse images from Movie S1 of a bilayer island exposed to BIN1-mEGFP. Membrane is colored in gray and inverted in contrast. Fluorescence (B) and scanning electron microscopic (C) images showing BIN1 tubules on the bilayer island after 10 mins. Membrane is colored in gray and inverted in contrast. Tubules are marked by white arrows. (D) Time-lapse images from Movie S2 showing coiling of BIN1 tubules. BIN1-mEGFP is colored in gray and inverted in contrast. (E) Time-lapse images from Movie S4 showing initiation and growth of BIN1 tubules, marked by white arrows. BIN1-mEGFP is colored in gray and inverted in contrast. (F) Plot showing growth rate of BIN1 tubules. Time is normalized to when distinct tubules because apparent. Data represent the mean ± SD of 9 tubules. (G) Plot showing the radius of BIN1 tubules. Separate uncoiled tubules were analyzed, and the data represent the mean ± SD of 37 tubules. (H) Time-lapse images from Movie S5 showing the formation of BIN1-mEGFP oligomers marked by white arrows on the bilayer island. Membrane is colored in gray and inverted in contrast. (I) Plot showing kinetics of BIN1 oligomerization and membrane tubulation fitted to a segmental linear regression equation. Data represent the mean ± SD of fluorescence intensities of 5 independent events.

We imaged the tubulation reaction at a high frame rate and analyzed tubules before they began to coil. The mild flow of buffer laid down emergent tubules, which allowed them to be easily imaged (Fig. 1E, Movie S4). Tubules grew at an apparent rate of ∼0.9 μm.s^-1^ (Fig. 1F). From a fluorescence-based calibration procedure, the estimated size of tubules was ∼17 nm in radius (Fig. 1G), which agrees well with the BIN1 scaffold dimension from cryo-EM reconstructions ^36^ and our previous estimates of BIN1 scaffolds on membrane nanotubes ^37^. To monitor early stages of the tubulation reaction, we flowed a low concentration of 50 nM BIN1 and imaged the island. The protein first bound uniformly and with time organized into foci, which likely represent oligomers, that diffused on the island (Fig. 1H, white arrows, Movie S5). BIN1 oligomers coincided with high membrane fluorescence because of tubulation of the underlying bilayer (Fig. 1H). Tracking fluorescence changes on these oligomers showed a sharp rise in both BIN1 and membrane fluorescence, signifying kinetics of nucleation and growth of a BIN1 tubule (Fig. 1I). Surprisingly, the onset of rise in BIN1 fluorescence occurred before the onset of rise in membrane fluorescence, by almost 20 s (Fig. 1I). This indicates that BIN1 first forms an oligomer that is likely planar in geometry, which grows by recruiting proteins from solution and eventually acquires curvature and the capacity to tubulate the membrane. Thus, BIN1 oligomerization and membrane tubulation are sequential and not concurrent processes.

### CNM-linked BIN1 mutants are defective in membrane binding and tubulation

The BIN1 N-BAR domain comprises of a trihelical coiled-coil domain that forms a crescent shaped dimer with positively charged residues on the concave, membrane binding surface (Fig. 2A) ^38^. Despite their structural similarity, BIN1 shows significant differences compared to other BAR domain-containing proteins ^36^. BIN1 tubules are more rigid because of additional interactions between neighboring BAR domains and tighter protein packing on the tubule ^36^. Cryo-EM reconstructions indicate that one tip (residues 130-190) of the N-BAR dimer wedges into the membrane while the other points outwards. Tip insertion into the membrane facilitates optimal packing of the oligomer ^36^. Like is the case for other N-BAR proteins, the unstructured N-terminal region forms an amphipathic H_0_ helix upon binding membranes and its insertion bends the membrane and stabilizes protein-protein contacts in the packed oligomer ^38–40^. Furthermore, tip residues have also been shown to become ordered upon membrane insertion ^39^. Importantly, mutations in the tip and the H_0_ helix residues are linked to CNM and previous analysis have shown defects in membrane binding and tubulation _10,21,41_. A more recently identified tip mutant R145C is also linked to CNM ^42^.

**Fig. 2.**
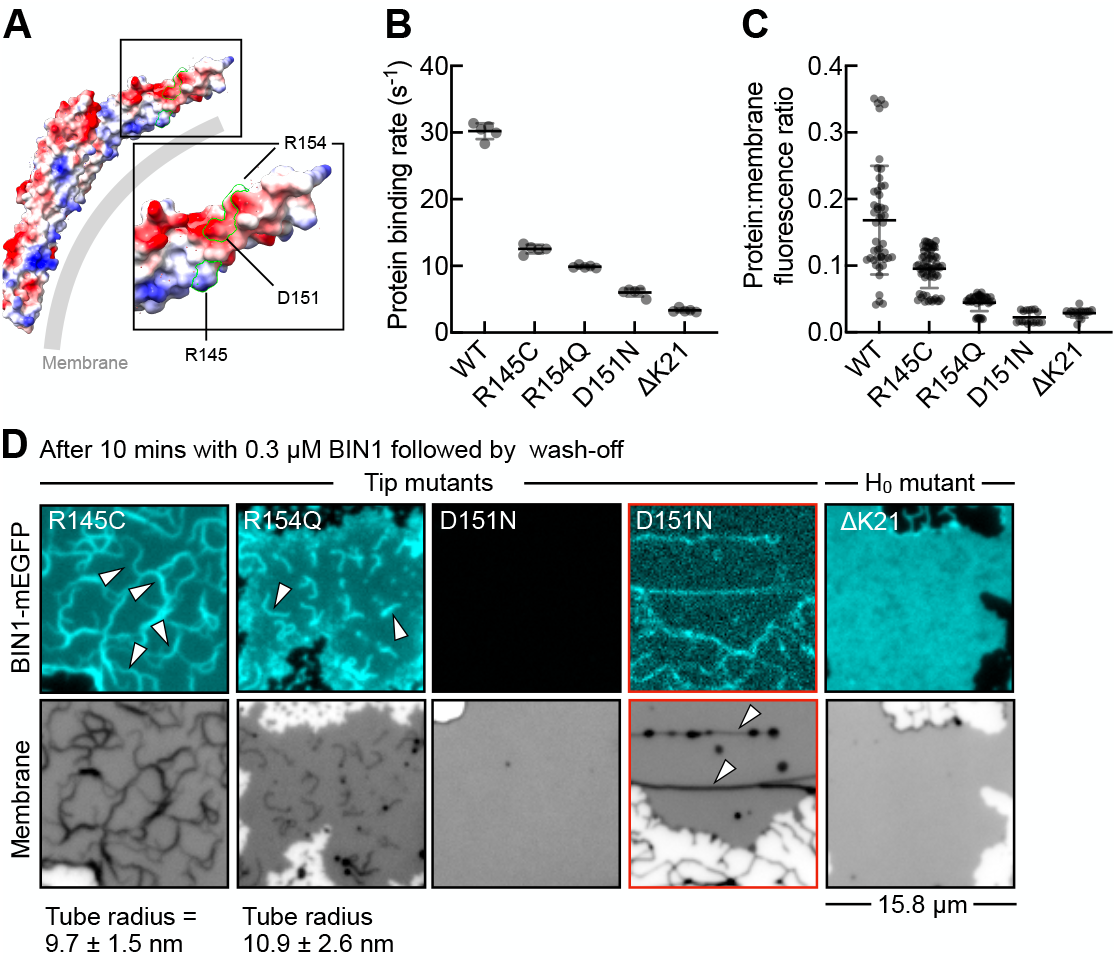
CNM-linked BIN1 mutants are defective in membrane binding and tubulation. (A) Structure of the BIN1 N-BAR domain (PDB: 2FIC) rendered as a space filling model and colored based on electrostatics using ChimeraX _56_. The structure marks the tip residues. (B) Plot showing the initial rates of membrane binding of BIN1 mutants. Data represent the mean ± SD of binding rates monitored in 5 different region-of-interest (ROI) placed on the bilayer island for each mutant. Binding rates were estimated by fitting the rise in BIN1-mEGFP fluorescence on bilayers to a segmental linear regression equation. (C) Membrane density of BIN1 mutants measured as the ratio of protein and membrane fluorescence after washing off excess protein. Data represent the mean ± SD of ratios from 48 ROIs for WT, 63 ROIs for R145C, 35 ROIs for R154Q, 17 ROIs for D151N and 17 ROIs for ΔK21 placed on the bilayer islands. (D) Fluorescence images showing the distribution of the indicated mutants on a bilayer island. Bilayers were incubated with 0.3 μM of proteins for 10 mins and the excess protein was washed off. Membrane is colored in gray and inverted in contrast. White arrows in R145C and R154Q panels mark membrane tubules. See Movies S6 and S7. Separate uncoiled tubules were analyzed, and the data represent the mean ± SD of 47 tubules for R145C and 33 tubules for R154Q. Red-outlined panels for D151N are bilayer islands with pre-existing membrane tubules marked by white arrows. The BIN1 fluorescence image has been contrast adjusted for visualization.

We tested the tip and H_0_ helix mutants by first analyzing the initial rates of membrane binding upon flowing proteins at a concentration of 0.1 μM. In comparison to WT, all the CNM-linked mutants displayed slower binding rates implying significant defects, but to different degrees (Fig. 2B). Indeed, analyzing protein density retained on the island after a buffer wash revealed that the mutants were severely defective in membrane binding (Fig. 2C). Raising the protein concentration to 0.3 μM and imaging the islands after a buffer wash showed retention of the tip mutants R145C and R154Q and associated tubulation (Fig. 2D). This indicates an oligomerization defect because tubulation can be partially rescued by increasing protein concentration. The R154Q mutant appeared to be more defective than R145C, both in the rates and extent of tubulation. Tubules formed with these mutants were spindly, with a radius of ∼10 nm for R145C and ∼11 nm for R154Q (Fig. 2D, Movie S6 and Movie S7), significantly thinner than that seen with WT (Fig. 1G) (P<0.001, Mann-Whitney’s test). Thus, the oligomerization residues reduce tubulation but surprisingly pack together into a forming a scaffold of thinner dimensions. In contrast, the other tip mutant D151N showed no retention on the bilayer (Fig. 2D), consistent with earlier results on PI(4,5)P_2_-containing membranes ^41^. We detected extremely low levels of D151N on pre-existing tubes on the island (Fig. 2D, white arrows in red-outlined panels), implying a defect in binding planar bilayers. Importantly, the H_0_ helix mutant ?K21 showed retention on the bilayer but no tubulation (Fig. 2D), thus emphasizing the important role of H_0_ helix insertion in membrane tubulation. The CNM-linked mutations therefore appear to cause distinct and separable effects on BIN1 functions in membrane binding, oligomerization and tubulation. D151N abrogates membrane binding, ?K21 can bind but is defective in curvature induction while the R145C and R154Q mutants reduce BIN1’s oligomerization. Together, these results provide important mechanistic insights into the pathway for BIN1-induced membrane tubulation. The ability to monitor binding kinetics and to distinguish binding from tubulation parses out subtle defects among the CNM-linked mutants and emphasizes the utility of bilayer islands.

### BIN1 and Dyn2 comprise a minimal two-component MTCF module

Besides membrane tubulation, BIN1 recruits dynamin during the formation of T-tubules in muscle cells ^20,21,25,43^. Flowing Dyn2 showed no effect because it barely bound the planar island, but it bound pre-existing buds and tubes (Fig. 3A, white arrows and Fig. 3B), consistent with its reported preference for binding membranes of high curvature ^37,44^. Bilayer islands therefore present a significant advantage because it would report Dyn2 functions only in response to membrane tubulation. To recreate a physiological scenario where both proteins encounter each other, we flowed a premixed solution of BIN1, Dyn2 and GTP onto the bilayer islands. Remarkably, this produced an outcome quite different from the extensive tubulation seen with BIN1 alone (Fig. 1A). The islands now displayed numerous bright foci (Fig. 3C, Movie S8), which contained both BIN1 and Dyn2. Importantly, bleaching the fluorescent lipid probe showed recovery on the underlying bilayer but not in foci (Fig. 3D, Movie S9), indicating that the foci are severed vesicles. We are unsure why the vesicles remain tethered to the island but multivalent interactions between BIN1 and Dyn2 on the vesicle and the bilayer could have contributed to the tethering. Together, these results indicate that BIN1 and Dyn2 comprise a minimal two-component module that can manage MTCF.

**Fig. 3.**
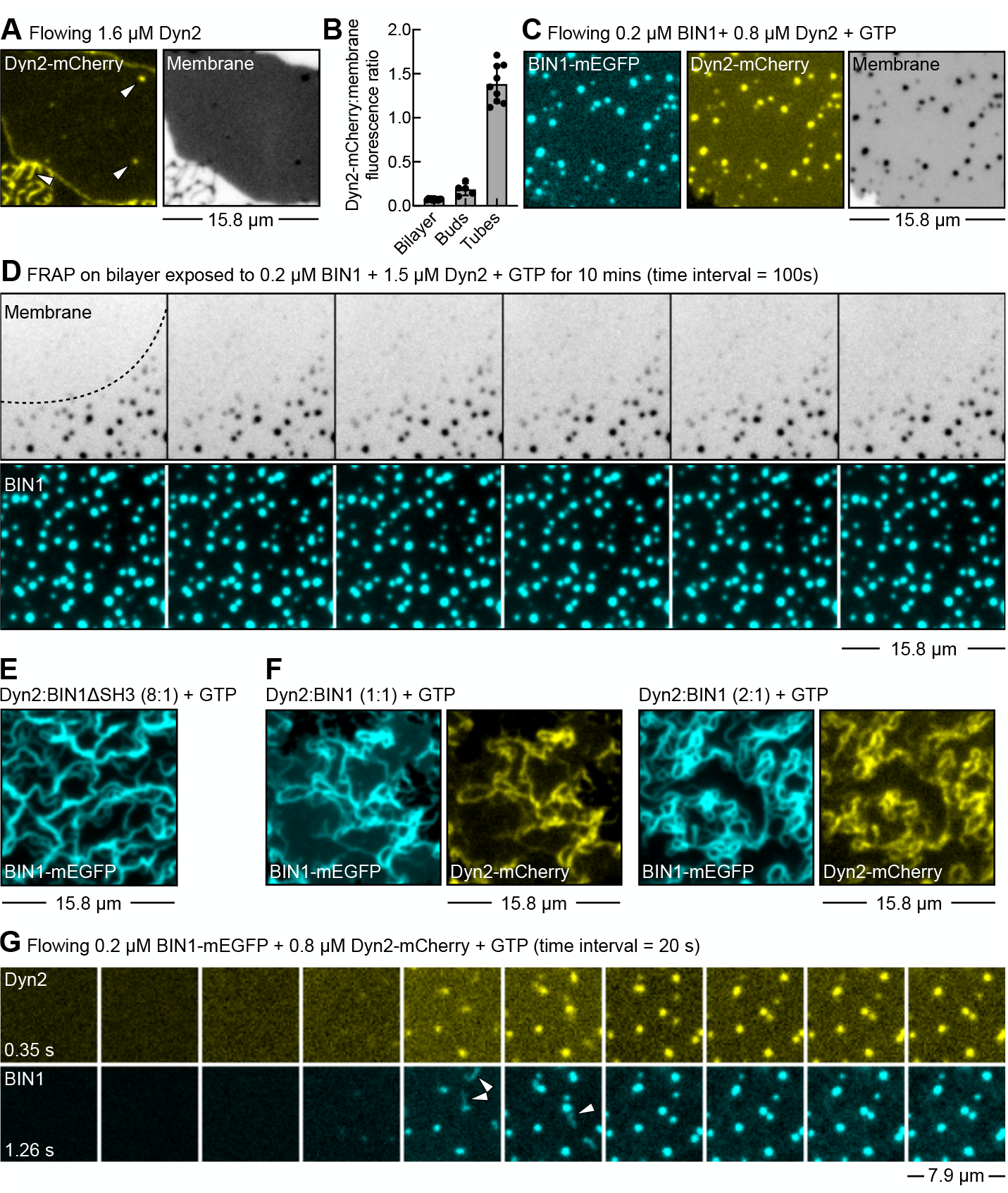
BIN1 and Dyn2 comprise a minimal two-component module that can manage MTCF. (A) Fluorescence image showing the distribution of Dyn2-mCherry on the bilayer island. Dyn2-mCherry is colored in yellow and membrane is colored in gray and inverted in contrast. White arrows mark pre-existing buds on the island. (B) Plot showing Dyn2 membrane density measured as the ratio of Dyn2-mCherry and membrane fluorescence on bilayers, buds and tubules. Data represent the mean ± SD of 11 bilayer patches, 5 buds and 9 tubes. (C) Fluorescence image of a bilayer island exposed to BIN1 with excess Dyn2 and GTP. Membrane is colored in gray and inverted in contrast. (D) Time-lapse images from Movie S9 showing recovery after bleaching the fluorescent lipid probe in a large area of the bilayer island exposed to BIN1 with Dyn2 and GTP. Dotted line represents the boundary of the bleached region. Membrane is colored in gray and inverted in contrast. (E) Fluorescence image showing the distribution of BIN1DSH3-mEGFP in the presence of excess Dyn2 with GTP. Bilayers were imaged after 10 mins incubation with 0.2 μM BIN1DSH3-mEGFP and 1.6 μM Dyn2 with 1 mM GTP. (F) Fluorescence image showing the distribution of BIN1-mEGFP in presence of Dyn2 with GTP. Bilayers were imaged after 10 mins of incubation with 0.2 μM BIN1-mEGFP and 0.2 μM or 0.4 μM Dyn2 with 1 mM GTP. (G) Time-lapse images from Movie S8 of bilayer islands exposed to BIN1-mEGFP and Dyn2-mCherry with 1 mM GTP. White arrows mark tubular intermediates.

Replacing BIN1 with a construct lacking the SH3 domain (BIN11ΔSH3) showed tubulation but no fission (Fig. 3E), indicating that Dyn2’s binding to BIN1 is necessary for fission. Furthermore, fission required a high relative concentration of Dyn2. Thus, reactions with equimolar concentrations of Dyn2 and BIN1 or even with a 2-fold molar excess of Dyn2 showed only tubulation and no fission (Fig. 3F). The binding affinity of the BIN1 SH3 domain for the Dyn2 PRD is ∼70 μM ^35^ and recent analysis reports an affinity of the BIN1 SH3 domain for Dyn2 to be ∼10 μM ^11^, which is still rather low. But Dyn2 was present on BIN1 tubules under conditions that showed no fission (Fig. 3F) or throughout reactions that led to the formation of vesicles (Fig. 3G, Movie S8). Therefore, the requirement of high relative concentrations of Dyn2 for fission is quite unexpected and cannot be attributed solely to the low binding affinity between these proteins.

### Pathway to MTCF

Dual channel time lapse imaging revealed the fleeting appearance of tubular intermediates that were positive for BIN1-mEGFP and Dyn2-mCherry (Fig. 3G, white arrows and Movie S8). To better analyze this reaction, we recorded time lapse images of BIN1-mEGFP mixed with excess Dyn2 at a high frame rate, for which imaging was restricted to a single channel. In the absence of GTP, flowing this mixture formed tubules from the bilayer island (Fig. 4A, black arrows, Movie S10). Tubules grew at a rate of ∼1 μm.s^-1^ (Fig. 4B), similar to that seen before with just BIN1 (Fig. 1F). Thus, the presence of Dyn2 does not affect the tubule growth rate. Furthermore, tubules grew while remaining tethered to the island (Fig. 4A, black arrows) and the fact that they grew at a rate like that seen in the absence of Dyn2 (Fig. 1F) suggests that the growth rate estimates reflect an intrinsic property of BIN1 oligomerization and are not influenced by tethering or buffer flow. Eventually, tubules started coiling upon themselves or with other tubules in the vicinity and coalesce into bright foci (Fig. 4A, green arrows). Flowing a mixture of BIN1 and Dyn2 in the presence of GTP led to the formation of tubules (Fig. 4C, black arrows), which quickly got severed (Fig. 4C, yellow arrows, Movie S11). Tubules sometimes displayed more than one fission event resulting in the formation of multiple tubule fragments. Like was seen in the absence of GTP, tubules showed a tendency to coil before or after fission and coalesce into bright foci (Fig. 4C, green arrows). Tubules grew at a rate of ∼0.8 μm.s^-1^ (Fig. 4D) and the time interval between the appearance and fission of a tubule for a number of such events was ∼5 s (Fig. 4E), which puts the combined fission rate at ∼0.2 s^-1^.

**Fig. 4.**
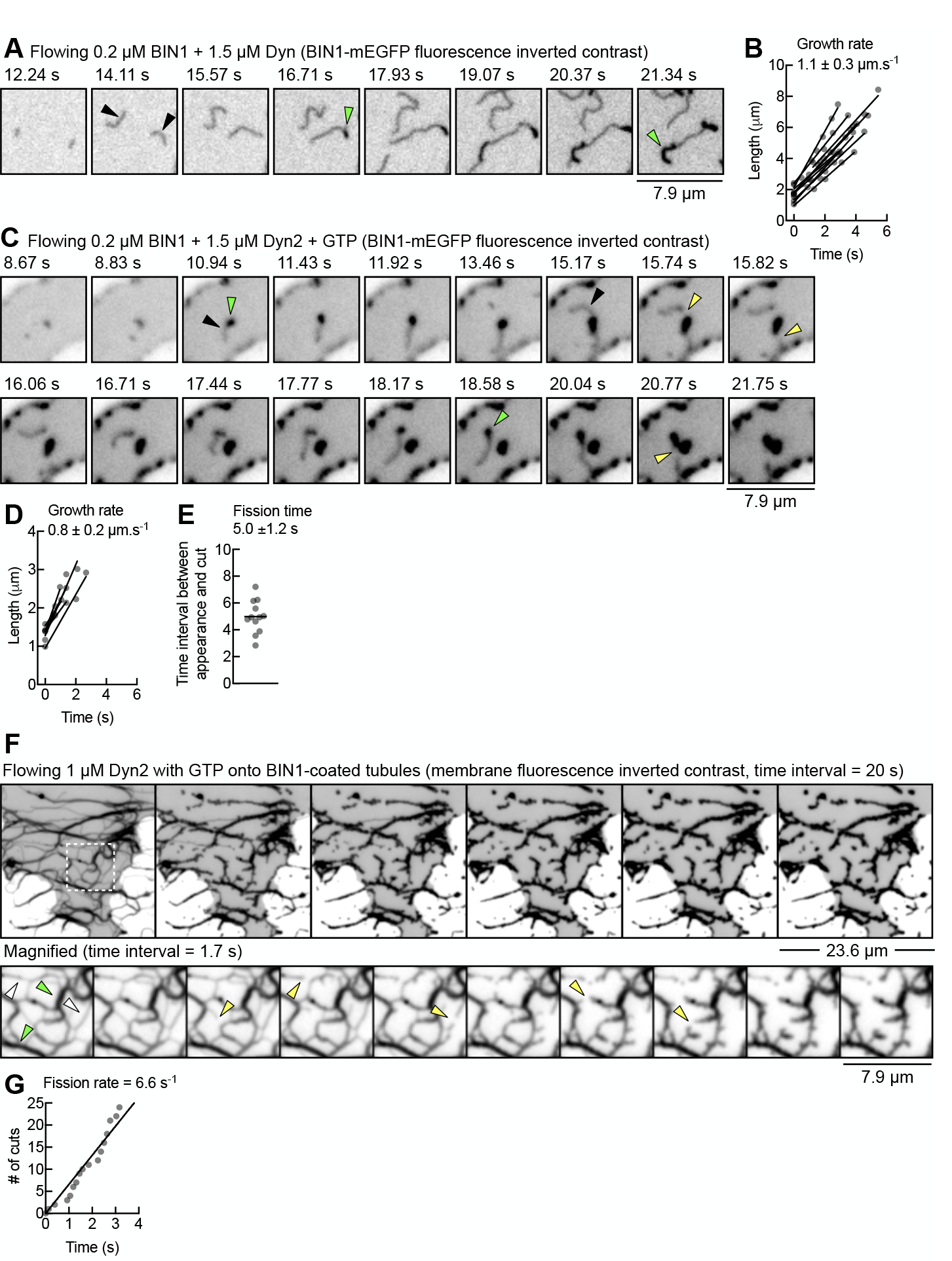
Pathway to MTCF. (A) Time-lapse images from Movie S10 of a bilayer island exposed to the indicated proteins in the absence of GTP. Black arrows mark single tubes and green arrows mark coiled regions. (B) Plot showing the growth rate of BIN1 tubules under conditions described in (A). Data represent the mean ± SD of 10 tubules. (C) Time-lapse images from Movie S11 of a bilayer island exposed to the indicated proteins in the presence of GTP. Black arrows mark single tubes, green arrows mark coiled regions and yellow arrows mark fission. (D) Plot showing the growth rate of BIN1 tubules under conditions described in (C). Data represent the mean ± SD of 6 tubules. (E) Plot showing the fission time, defined as the time interval between appearance and fission of a tubule. Data represent the mean ± SD of 12 events. (F) Time-lapse images from Movie S12 of preformed BIN1 tubules exposed to Dyn2 with GTP. Bottom panel is a magnified view of the ROI marked with dotted line in the top panel. White arrows mark single tubules, green arrows mark coiled tubules and yellow arrows mark fission. (G) Plot showing the cumulative frequency of cuts seen in (F) and fitted to a linear regression. The slope of this fit gives the fission rate.

The above-described reactions were carried out with premixed BIN1 and Dyn2, where Dyn2 appeared to be bound to BIN1 throughout the reaction (Fig. 3F, G). We wondered how this would compare to a sequential reaction, where Dyn2 was added to preformed BIN1 tubules. Bilayer islands were first incubated with BIN1, which formed a network of tubules, after which we imaged the network while flowing an excess of 1 μM Dyn2 with GTP. Dyn2 arrival caused fission of BIN1 tubules (Fig. 4F, Movie S12). But a closer look revealed that the fission events were largely restricted to tubes that were uncoiled (Fig. 4F, magnified panel, white arrows). Coiled tubules (Fig. 4F, green arrows) rarely got severed, thereby rendering fission to appear less processive under these conditions. Consequently, these reactions left behind long BIN1 tubule fragments. But plotting the cumulative frequency of cuts on uncoiled tubes gave us an apparent fission rate of ∼7 s^-1^ (Fig. 4G), which is considerably faster than the fission rate observed with premixed Dyn2 and BIN1 (Fig. 4E).

### Mechanistic basis for MTCF

Our results suggest that membrane tubulation and fission are temporally linked, i.e., tubules grow for a while before fission is apparent. This could merely arise from Dyn2’s preference for binding high curvature tubules such that fission manifests only upon the formation of a BIN1 tubule. But results with BIN11ΔSH3 indicate that the formation of a tubule of high curvature is by itself not sufficient for fission. Furthermore, Dyn2 flowed on to preformed BIN1 tubules shows a fission rate of ∼7 s^-1^ (Fig. 4F) while Dyn2 premixed with BIN1 shows a fission rate of 0.2 s^-1^, which consequently allows tubules to grow to a considerable length (Fig. 4C). This indicates that the Dyn2 incorporated in growing BIN1 tubules must be inhibited in fission and the requirement for high relative concentrations of Dyn2 could reflect the necessity to overcome this inhibition.

The obligatory requirement of BIN1 for tubulation makes it difficult to analyze the effects of BIN1 on Dyn2 functions. We therefore turned to analyzing Dyn2 functions on preformed nanotubes of the same lipid composition as bilayer islands. Control experiments revealed that BIN1 facilitates recruitment of Dyn2 onto nanotubes of a wide range of tube sizes (Fig. S2A, B). Flowing 0.1 μM Dyn2 with GTP caused rapid and processive severing of nanotubes (Movie S13). We then analyzed the effect of flowing 0.1 μM Dyn2 and GTP mixed with increasing concentrations of BIN1. Imaging the nanotubes after 10 mins revealed fission till a concentration of 0.6 μM BIN1 (Fig. 5A). At 0.6 μM BIN1, nanotubes were completely refractory to fission (Movie S14). This is apparent from estimates of the fraction of nanotubes showing at least one cut after 10 mins (Fig. 5B). Time-lapse imaging revealed inhibitory effects even at lower BIN1 concentrations. This is apparent from estimates of the frequency of cuts on individual nanotubes after a short 2 min interval (Fig. 5C). BIN1 therefore inhibits Dyn2-catalyzed membrane fission in a dose-dependent manner.

**Fig. 5.**
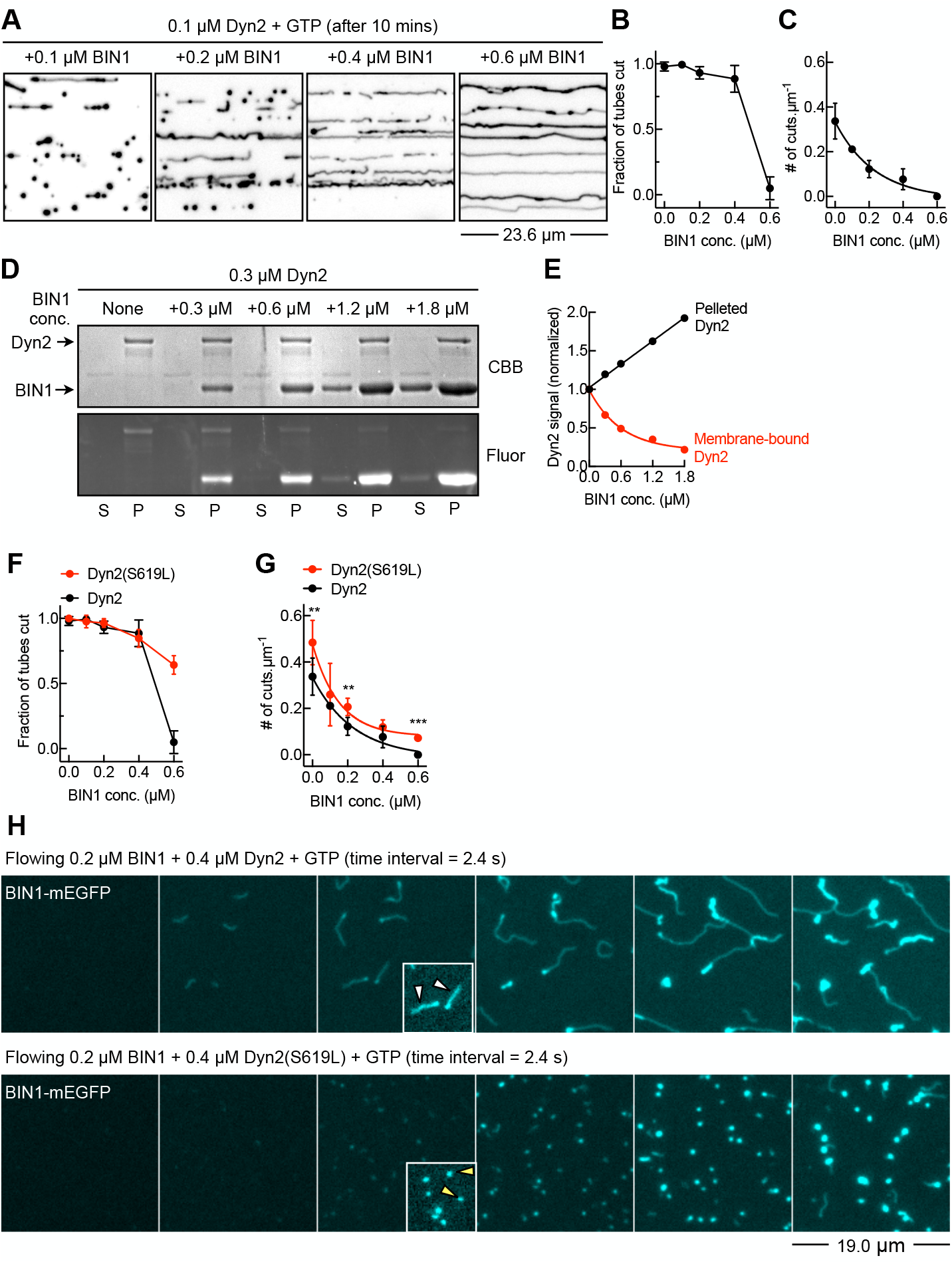
Mechanistic basis for MTCF. (A) Fluorescence images of membrane nanotubes after being exposed to the indicated proteins for 10 mins. Membrane is colored in gray and inverted in contrast. (B) Plot showing the fraction of nanotubes showing at least one cut in presence of 0.1 μM Dyn2 and 1 mM GTP with increasing concentrations of BIN1. Data represent the mean ± SD from 6 separate fields of nanotubes. (C) Plot showing the frequency of cuts on single nanotubes after 2 mins of exposure to 0.1 μM Dyn2 and 1 mM GTP with increasing concentrations of BIN1. Data represent the mean ± SD from 7-10 nanotubes. (D) Results from a coupled liposome co-sedimentation and PLiMAP experiment showing in-gel fluorescence (Fluor) and Coomassie brilliant blue (CBB) staining of Dyn2 and BIN1 in the supernatant (S) and pellet (P) fractions. (E) Plot showing densitometric (black) and fluorescence (red) quantitation of Dyn2 in the pellet fraction in (D). (F) Plot showing the fraction of nanotubes showing at least one cut in presence of 0.1 μM Dyn2(S619L) and 1 mM GTP with increasing concentrations of BIN1. Data represent the mean ± SD from 6 separate fields of nanotubes. Data for Dyn2 is reproduced from (B) for comparison. (G) Plot showing the frequency of cuts on single nanotubes after 2 mins of exposure to 0.1 μM Dyn2(S619L) and 1 mM GTP with increasing concentrations of BIN1. Data represent the mean ± SD from 5-10 nanotubes. Data for Dyn2 is reproduced from (C) for comparison. Statistical significance was estimated using Mann-Whitney’s test where *** denotes P =0.003 and ** denotes P = 0.0015. (H) Time-lapse images from Movie S15 showing the bilayer island exposed to the indicated concentrations of proteins with 1 mM GTP. Insets show magnified images of intermediates adjusted in contrast for clarity. White arrows mark tubules and yellow arrows mark vesicles.

The N-BAR and SH3 domain containing protein endophilin inhibits Dyn1 functions by interfering with the formation of Dyn1 scaffolds, necessary for stimulated GTP hydrolysis and fission ^45^. But a mechanism involving structural inhibition should have influenced BIN1 tubule growth rates in the presence of Dyn2, especially when they are both present on the tubule, which is not the case (Fig. 1F, 4B). How then can the inhibition of Dyn2 functions by BIN1 be explained? To address this, we performed a coupled liposome co-sedimentation and PLiMAP assay with Dyn2 and BIN1. Co-sedimentation reports on the total levels of liposome-associated proteins while PLiMAP reports on their proximity to the membrane interface, which is a more stringent measure of membrane binding. Liposomes were incubated with 0.3 μM Dyn2 mixed with a range of BIN1 concentrations and incubated for 30 mins. The mixture was then exposed to UV and sedimented. In the absence of BIN1, a substantial fraction of Dyn2 co-sediments with liposomes (Fig. 5D, CBB panel), and is also fluorescent (Fig. 5D, fluor panel), indicating that readouts from co-sedimentation and PLiMAP are consistent with each other. Increasing BIN1 concentrations does not substantially alter Dyn2 levels in the pellet and a densitometric analysis in fact reveals an increase in Dyn2 levels (Fig. 5D, E). This is consistent with BIN1 facilitating recruitment of Dyn2 on nanotubes (Fig. S2). But surprisingly, the fluorescence signal of Dyn2 in the pellet shows a significant decline, with a 75% reduction at a 1:6 Dyn2:BIN1 ratio (Fig. 5D, E). Thus, increasing BIN1 levels recruits more Dyn2 but lowers the fraction of membrane bound Dyn2. This could explain the inhibition in fission seen with higher BIN1 concentrations (Fig. 5B, C). Since the inhibition is dose-dependent, increasing Dyn2 concentration would overcome this inhibition and explains why fission requires a high relative concentration of Dyn2. Tubules formed with BIN11ΔSH3 are resilient to fission, signifying that the SH3-PRD interaction helps overcome the inhibition likely because the multivalent SH3-PRD interaction would help attain the critical Dyn2 concentration required for fission on the tubule at a lower bulk Dyn2 concentration.

Dyn2 binds PI(4,5)P_2_ in the membrane through the pleckstrin-homology domain (PHD). Our results indicate that BIN1 inhibits Dyn2 functions by reducing its membrane binding. The PHD binds the stalk domain and occludes an oligomerization interface thereby rendering Dyn2 autoinhibited in solution ^46^. The stalk domain mutation S619L interferes with this binding and relieves autoinhibition thereby facilitating Dyn2 oligomerization. But this mutation also enhances Dyn2’s membrane binding and facilitates fission ^25,43,47^. Importantly, the S619L represents a CNM-linked gain-of-function mutation ^18,48,49^. On membrane nanotubes, increasing concentrations of BIN1 only partially inhibited the Dyn2(S619L)-catalyzed fission. Thus, in the presence of 0.6 μM BIN1, while Dyn2 showed no fission, the fraction of severed tubes with Dyn2(S619L) rose to ∼0.75 (Fig. 5F). This effect is more apparent on the frequency of cuts on individual nanotubes after 2 mins of the fission reaction. Dyn2(S619L) showed a significantly higher fission efficiency than Dyn2 (Fig. 5G) and was less susceptible to BIN1-mediated inhibition than Dyn2 (Fig. 5G). Finally, we tested Dyn2(S619L) mixed with BIN1 on bilayer islands. As shown earlier, reactions with 2-fold molar excess of Dyn2 over BIN1 showed only tubulation and no fission (Fig. 3F). This is apparent in time-lapse images of the bilayer island exposed to 0.2 μM BIN1 and 0.4 μM Dyn2 with GTP (Fig. 5H, Movie S15). But reactions with 0.2 μM BIN1 and 0.4 μM Dyn2(S619L) with GTP showed remnants of severed tubules, which appeared as foci, indicative of fission (Fig. 5H, Movie S15). The PI stretch in BIN1 binds and clusters PI(4,5)P_2 24_, and a likely mechanism by which BIN1 inhibits Dyn2 function is by sequestering PI(4,5)P_2_. This is consistent with observations that the inhibitory effect of BIN1 on Dyn2 can be overcome by the S619L mutation, which confers Dyn2 with a higher membrane binding affinity.

Together, our results from the use of cushioned bilayer islands reveal that BIN1 and Dyn2 together constitute a minimal two-component MTCF module. Importantly, CNM-linked mutants of BIN1 reduce membrane tubulation while those linked to Dyn2 enhance fission, thereby impacting MTCF functions of this module.

## Discussion

Here, we establish a facile template and a broadly applicable workflow for analyzing dynamics of membrane tubulation. Our results provide novel insights into the functional partnership between BIN1 and Dyn2. CNM-linked BIN1 and Dyn2 mutants tested here display aberrant functions in membrane tubulation and fission and these results become relevant in understanding CNM-like pathology. To the best of our knowledge, these are the first results that reconstitute and thereby reveal dynamics of a minimal MTCF reaction, from which emerge key regulatory principles.

The architecture of stable T-tubules in tissues has been well studied but an understanding of the early steps in their biogenesis is only beginning to emerge. Live imaging of T-tubule development in zebrafish embryos suggests a pathway wherein BIN1 along with other proteins form dynamic endocytic tubules at the plasma membrane, some of which are captured and stabilized by contacts with the sarcoplasmic reticulum ^5^. This endocytic capture model highlights similarities between early stages of T-tubule biogenesis and clathrin-independent trafficking pathways that involve a tubular membrane intermediate. Recent work visualizing the plasma membrane of unroofed differentiated myotubes shows BIN1-positive tubules emanating from ring-like platforms composed of caveolar proteins ^50^. Dyn2 is a stable component of developing and mature T-tubules but surprisingly also functions to sever BIN1 tubules and a dysregulation of this process is linked to CNM ^17,18,20,21,25,43^. Our results demonstrate that the relative levels of Dyn2 and BIN1 determine whether tubules remain intact or get severed. BIN1 competitively inhibits Dyn2 functions at the level of its membrane binding. At low ratios of Dyn2 and BIN1, this inhibition prevents severing of BIN1 tubules. High Dyn2 concentrations overcome this inhibition and renders the BIN1 tubule susceptible to fission. BIN1 and Dyn2 levels are mutually regulated during muscle development ^16^. Perhaps, at early stages during development where Dyn2 levels are relatively high, fission of emergent BIN tubules acts as a quality control mechanism. Tubules that form contacts with the sarcoplasmic reticulum are left intact while others are pruned and turned over. Hypoactive BIN1 mutants would produce fewer and structurally defective tubules while hyperactive Dyn2 mutants would result in excessive pruning, both of which would lower tubule densities and eventually impact T-tubule architecture.

Our results provide insights into mechanisms that regulated MTCF, which could be relevant in vesicular transport. MTCF dynamics seen with BIN1 and Dyn2, where the tubule extends for a considerable length before undergoing fission, is reminiscent of several intracellular transport reactions involving a tubular intermediate^2,3,7^. Pathways that form vesicular transport carriers (VTCs), such as during clathrin-mediated endocytosis where the coat imposes a bud-like architecture on emergent transport intermediates are quite well-characterized in terms of mechanisms that regulate their budding and fission. The budding process defines a neck, which is the region with the highest membrane curvature, and drives the assembly of BAR domain containing proteins that eventually recruit dynamin for fission ^13^. On the other hand, mechanisms that coordinate growth and fission of tubular transport carriers (TTCs) remain less understood ^51^. TTCs are apparent in numerous clathrin-independent endocytic and endocytic recycling pathways ^2,3,7,52–54^ For a TTC-based transport pathway to function optimally, membrane tubulation must be coordinated with fission. Excessive tubulation or premature fission would affect throughput of the transport reaction. Furthermore, tubules display the same membrane curvature along their entire length, so a mechanism that utilizes a gradient of curvature to define the fission site is untenable. Proteins such as sorting nexins (SNXs), which contain a membrane binding and bending domain are overrepresented in intracellular TTC-based transport pathways. SNXs contain a PX domain that binds phosphoinositide lipids and a BAR domain that forms a tubular scaffold or coat ^54,55^. The presence of these domains would render SNXs capable of both initiating and progressively tubulating donor membranes. But our results suggest that the same features could also intrinsically regulate fission. Our results with BIN1, which also contains a membrane binding PI stretch and membrane bending BAR domain, informs of a mechanism that regulates MTCF. BIN1 recruits Dyn2 through SH3-PRD interactions but inhibits its membrane binding, likely because the PI stretch of BIN1 competes with the PHD of Dyn2 for PI(4,5)P_2_ binding. This dual effect of potentiating recruitment but inhibiting membrane binding signifies a novel mechanism for how a protein scaffold regulates MTCF. While the exact mechanism that manages release of TTCs remains unclear, a similar competition for phosphoinositide lipids between the SNX coat and the putative fission catalyst could determine fission rates, thereby regulating the throughput of TTC-based transport pathways. Reconstituting such reactions using the membrane template described here represents an exciting avenue for future research.

## Acknowledgments

SB thanks the Indian Institute of Science Education and Research, Pune for a graduate fellowship. TJP thanks the Howard Hughes Medical Institute for funding support. We thank Girish Deshpande and Pucadyil Lab members for comments on the manuscript.

## Author contributions

SB and TJP designed research; SB performed research; SB contributed new reagents; SB and TJP analyzed data; and SB and TJP wrote the paper.

## Methods

### Expression, purification, and fluorescent labelling of proteins

Human BIN1-EGFP (isoform 8) (Addgene plasmid #27305) and human dynamin2 C-terminally tagged to mCherry were cloned in pET15b with an N-terminal 6xHis and a C-terminal StrepII tag. Human dynamin 2 was cloned in pET15B with a C-terminal StrepII tag. Mutations were introduced using PCR. All clones were confirmed by sequencing. BL21(DE3) cells were grown at 37 °C to an OD_600_ of 0.6. Protein expression was induced with 0.1 mM IPTG, and the cells were transferred to 18 °C and left for 16 h with shaking. Bacterial cells were pelleted and stored at -40 °C. The frozen bacterial pellet was thawed in 20 mM HEPES pH 7.4 with 500 mM NaCl, 1% Triton X-100 and a protease inhibitor cocktail tablet (Roche). Cells were lysed by sonication in an ice-water bath. The lysate was spun at 30,000 g for 20 min. For proteins containing 6xHis and StrepII tags (BIN1, BIN1-EGFP, dynamin2-mCherry), the supernatant was incubated with HisPur™ Cobalt Resin (Thermo Fisher Scientific). The resin was washed with 20 mM HEPES pH 7.4 with 500 mM NaCl, and the bound protein was eluted with 20 mM HEPES pH 7.4 with 500 mM NaCl and 250 mM imidazole. The elution was loaded onto a StrepTrap HP column (GE Lifesciences), washed with 20 mM HEPES pH 7.4 with 150 mM NaCl and eluted with the same buffer containing 2.5 mM desthiobiotin. For dynamin2, the supernatant was directly loaded onto a StrepTrap HP column, washed with 20 mM HEPES pH 7.4 with 500 mM NaCl, exchanged for 20 mM HEPES pH 7.4 with 150 mM NaCl and eluted with the same buffer containing 2.5 mM desthiobiotin. Post binding to the StrepTrap HP column and buffer exchange, dynamins were additionally washed with buffer containing 100 mM EDTA to remove bound nucleotides prior to elution. Proteins were spun at 100,000 g to remove aggregates before use in assays. Protein concentrations were estimated using a BCA Protein Assay Kit (Takara). For FRAP experiments, BIN1 was labeled with 5-fold excess of Texas Red-C5-maleimide (Invitrogen) in 20 mM HEPES pH 7.4 with 150 mM NaCl. The reaction was quenched with excess DTT and the unreacted dye was removed by extensive dialysis.

### Proximity-based labeling of membrane-associated proteins (PLiMAP)

1,2-dioleoyl-sn-glycero-3-phosphocholine (DOPC), 1,2-dioleoyl-sn-glycero-3-phospho-L-serine (sodium salt), (DOPS), 1,2-dioleoyl-sn-glycero-3-phospho-(1′-myo-inositol) derivatives of phosphoinositides and BODIPY FL or BODIPY TMR phosphatidylethanolamine were from Avanti Polar Lipids. Diazirine derivatives of BODIPY FL or BODIPY TMR phosphatidylethanolamine were generated as described earlier ^33,34^. PLiMAP assays were carried out as described earlier ^33,34^. Briefly, lipids were aliquoted at desired ratios along with 1 mol% of BODIPY FL or BODIPY TMR-diazirine phosphatidylethanolamine in a glass tube and dried under high vacuum for 30 min. Deionized water was added to the dried lipids to achieve a final concentration of 1 mM. Lipids were hydrated at 50 °C for 30 min, vortexed vigorously and extruded through 50 nm (for BIN1 or Dyn2 alone) or 200 nm (for BIN1 and Dyn2 together) pore-size polycarbonate filters (Whatman). 100 μM liposomes was incubated with the indicated concentrations of proteins in a final volume of 30 μL. The reaction was incubated in the dark at room temperature for 30 min and exposed to 365 nm UV light for 1 min of an intensity of 200 mJ.cm^-2^ (UVP crosslinker CL-1000L). For the coupled liposome co-sedimentation and PLiMAP assay, the mixture was spun at 100,000 g. Samples were boiled and resolved using SDS-PAGE. Gels were first imaged for BODIPY fluorescence on a Typhoon Biomolecular imager (Amersham) and later fixed and stained with Coomassie Brilliant Blue (CBB). Binding data were fitted to a one-site binding isotherm using Graphpad Prism.

### Field emission scanning electron microscopy

BIN1 tubles were fixed inside the flow chamber. Samples were fixed with glutaraldehyde (3% w/v) for 10 mins and rinsed with 1% SDS and then with excess water. Samples were dehydrated by sequentially flowing 10, 20, 40, 60, 80 and 100% ethanol. Fluorescence imaging at each step confirmed no gross changes to protein distribution during sample preparation. The chamber was disassembled, and coverslips were kept under vacuum overnight. Samples were gold-coated using a Q150T turbo-pumped sputter coater (Quorum Technologies) and imaged on an Ultra Plus field emission scanning electron microscope (Zeiss) using a 1.9-kV electron beam and secondary electron detector.

### PEG-cushioned bilayer islands and membrane nanotubes

Lipids were aliquoted at desired ratios to a final concentration of 1 mM in chloroform. Lipid mixes also contained 0.5 mol% of the fluorescent lipid *p*Texas-Red DHPE (Thermo Fisher Scientific). For photobleaching experiments, lipid mixes contained 0.5% of BODIPY PE (Avanti Polar). 3 μl of the lipid mix was spread as a band with a glass syringe on glass coverslips covalently conjugated with PEG400 or PEG8000 ^32^, dried, and assembled inside an FCS2 flow chamber (Bioptechs), which was connected to a peristaltic pump (Bioptechs). The lipid film was hydrated by flowing buffer at a low flow rate, which washed off excess membranes and left behind bilayer islands at the region where the lipid was spread and membrane nanotubes downstream of the island. The chamber was pre-equilibrated with an oxygen scavenger cocktail ^32^ in 20 mM HEPES pH 7.4 with 150 mM NaCl and time-lapse images were acquired while flowing desired proteins in the same buffer in the absence or presence of 1 mM GTP (Jena Bioscience) and 1 mM MgCl_2_.

### Fluorescence imaging and image analysis

Reactions were imaged through 100x, 1.4 NA oil-immersion objective on an Olympus IX83 inverted microscope connected to an LED light source (CoolLED) and an Evolve 512 EMCCD camera (Photometrics). Image acquisition was controlled by μManager and images were analyzed using Fiji ^57^. Area photobleaching experiments were carried out in a wide-field microscopy mode. Bilayer islands were first incubated with the desired proteins. Excess protein was washed off and the lipid probe in the bilayer island was bleached by increasing the incident light intensity. This bleached a large area of the bilayer. The incident light intensity was lowered, and the bleached region was imaged at 10 or 20 s intervals to monitor recovery of fluorescence. Tube sizes were estimated based on a calibration procedure as described earlier ^32^. Fission efficiency was calculated by estimating the fraction of tubes that showed at least one cut after 10 mins or by estimating the number of cuts per tube after 2 min from time-lapse movies.

## Supplementary movie legends

**Movie S1.** BIN1-induced membrane tubulation. Movie shows the bilayer island responding to flowing 0.2 μM BIN1-mEGFP. BIN-mEGFP is colored cyan and membrane is colored gray and inverted in contrast. The field of view is 31.5 x 31.5 μm. Images were acquired at a time interval of 2.45 s and the movie is played at 5 fps.

**Movie S2.** Coiling of BIN1 tubules. Movie shows BIN1 tubules imaged after 10 mins. BIN-mEGFP is colored cyan. The field of view is 31.5 x 31.5 μm. Images were acquired at a time interval of 1.3 s and the movie is played at 5 fps.

**Movie S3.** Fluorescence recovery after photobleaching the lipid probe on the bilayer island exposed to BIN1. BIN-mEGFP is colored cyan and membrane is colored gray and inverted in contrast. The field of view is 15.8 x 15.8 μm. Images were acquired with a time interval of 10 s and the movie is played at 5 fps.

**Movie S4.** Early stages of BIN1-induced membrane tubulation. Movie shows a bilayer island responding to flowing 0.2 μM BIN1 acquired at a high frame rate in the membrane fluorescence channel. Membrane fluorescence is colored gray and inverted in contrast. The field of view is 15.8 x 15.8 μm. Images were acquired with a time interval of 0.132 s and the movie is played at 5 fps.

**Movie S5.** BIN1 oligomerization and membrane tubulation. Movie shows a bilayer island responding to flowing 50 nM BIN1-mEGFP. BIN-mEGFP fluorescence is colored cyan and membrane is colored gray and inverted in contrast. The field of view is 15.8 x 15.8 μm. Images were acquired at a time interval of 2.45 s and the movie is played at 5 fps.

**Movie S6.** Membrane tubulation with BIN1(R145C). Movie shows a bilayer island responding to flowing 0.3 μM BIN1(R145C). BIN-mEGFP is colored cyan and membrane is colored gray and inverted in contrast. The field of view is 15.8 x 15.8 μm. Images were acquired at a time interval of 2.45 s and the movie is played at 5 fps.

**Movie S7.** Membrane tubulation with BIN1(R154Q). Movie shows a bilayer island responding to flowing 0.3 μM BIN1(R154Q). BIN-mEGFP is colored cyan and membrane is colored gray and inverted in contrast. The field of view is 15.8 x 15.8 μm. Images were acquired at a time interval of 2.45 s and the movie is played at 5 fps.

**Movie S8.** Tubulation coupled with fission. Movie shows a bilayer island responding to flowing 0.2 μM BIN1-mEGFP and 0.8 μM Dyn2-mCherry with 1 mM GTP. BIN-mEGFP is colored cyan and Dyn2-mCherry is colored yellow. The field of view is 7.9 x 7.9 μm. Images were acquired with at a time interval of 2.45 s and the movie is played at 5 fps.

**Movie S9.** Fluorescence recovery after photobleaching the lipid probe on the bilayer island exposed to BIN1 and Dyn2 with GTP. BIN-mEGFP fluorescence is colored in cyan and membrane is colored in gray and inverted in contrast. 15.8 x 15.8 μm. Images were acquired with a time interval of 20 s and the movie is played at 5 fps.

**Movie S10.** High speed imaging of membrane tubulation. Movie shows a bilayer island responding to flowing 0.2 μM BIN1-mEGFP and 0.8 μM Dyn2 acquired at a high frame rate. BIN-mEGFP is colored in gray and inverted in contrast. The field of view is 15.8 x 15.8 μm. Images were acquired at a time interval of 80 ms and the movie is played at 5 fps.

**Movie S11.** High speed imaging of membrane tubulation coupled with fission. Movie shows a bilayer island responding to flowing 0.2 μM BIN1-mEGFP and 1.5 μM Dyn2 with 1 mM GTP. BIN-mEGFP is colored in gray and inverted in contrast. The field of view is 15.8 x 15.8 μm. Images were acquired at a time interval of 80 ms and the movie is played at 5 fps.

**Movie S12.** Fission of preformed BIN1 tubules. Movie shows a network of BIN1 tubules responding to flowing 1 μM Dyn2 with 1 mM GTP. Membrane is colored gray and inverted in contrast. The field of view is 23.6 x 23.6 μm. Images were acquired at a time interval of 132 ms and the movie is played at 5 fps.

**Movie S13.** yn2-catalyzed fission of nanotubes. Movie shows nanotubes responding to flowing 0.1 μM Dyn2 with 1 mM GTP. Membrane is colored gray and inverted in contrast. The field of view is 23.6 x 23.6 μm. Images were acquired at a time interval of 2.45 s and the movie is played at 5 fps.

**Movie S14.** Inhibition of Dyn2-catalyzed fission of nanotubes by BIN1. Movie shows membrane nanotubes responding to flowing 0.1 μM Dyn2 mixed with 0.6 μM BIN1 and 1 mM GTP. Membrane is colored gray and inverted in contrast. The field of view is 23.6 x 23.6 μm. Images were acquired at a time interval of 2.45 s and the movie is played at 5 fps.

**Movie S15.** Comparison between Dyn2 and Dyn2(S619L). Movie shows a bilayer island responding to flowing 0.2 μM BIN1-mEGFP with 0.4 μM Dyn2 (left) or 0.2 μM Dyn2(S619L) (right) with 1 mM GTP. BIN-mEGFP is colored cyan. The field of view is 18.9 x 18.9 μm. Images were acquired at a time interval of 2.45 s and the movie is played at 5 fps.

---

**Fig. S1.**
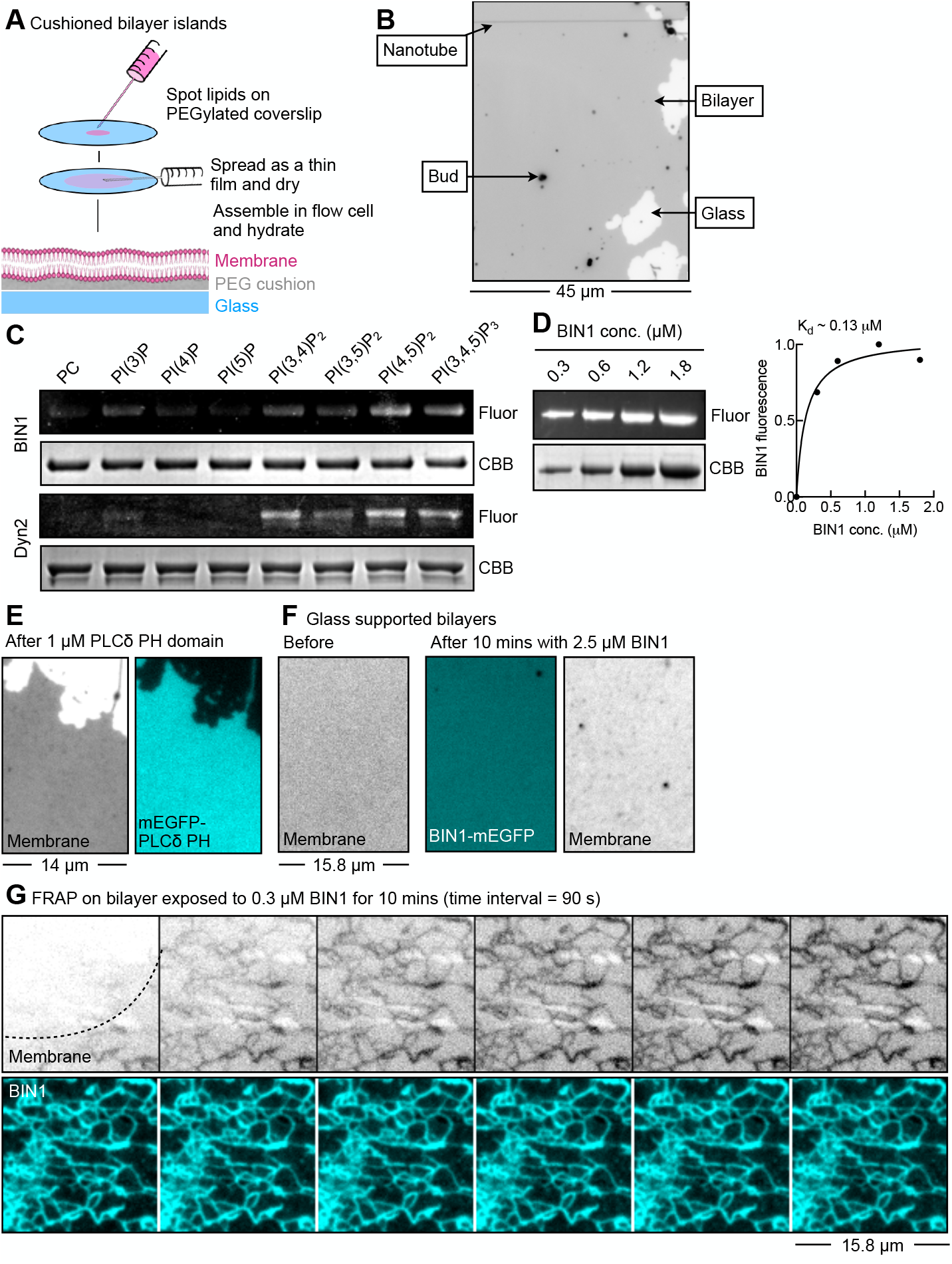
(A) Schematic showing the preparation of cushioned bilayer islands. (B) Fluorescence image of a bilayer island with membrane buds and nanotubes. Membrane is colored in gray and inverted in contrast. (C) Results from a PLiMAP experiment showing in-gel fluorescence (Fluor) and Coomassie brilliant blue (CBB) staining of BIN1 and Dyn2 on liposomes containing various phosphoinositide lipids. (D) Results from a PLiMAP experiment showing in-gel fluorescence (Fluor) and Coomassie brilliant blue (CBB) staining of BIN1. BIN1 fluorescence is plotted against BIN1 concentration and fitted to a one-site binding isotherm to estimate the binding affinity (K_d_). (E) Fluorescence images showing a bilayer island incubated with the PI(4,5)P_2_ sensor mEGFP-PLCd PH domain. Membrane is colored in gray and inverted in contrast. (F) Fluorescence images showing a glass-supported lipid bilayer before and after exposure to BIN1-mEGFP. Membrane is colored gray and inverted in contrast. (G) Time-lapse images from Movie S3 showing recovery after photobleaching the fluorescent lipid probe in a large area of the bilayer island displaying BIN1 tubules. Dotted line represents the boundary of the bleached region. Membrane is colored in gray and inverted in contrast.

**Fig. S2.**
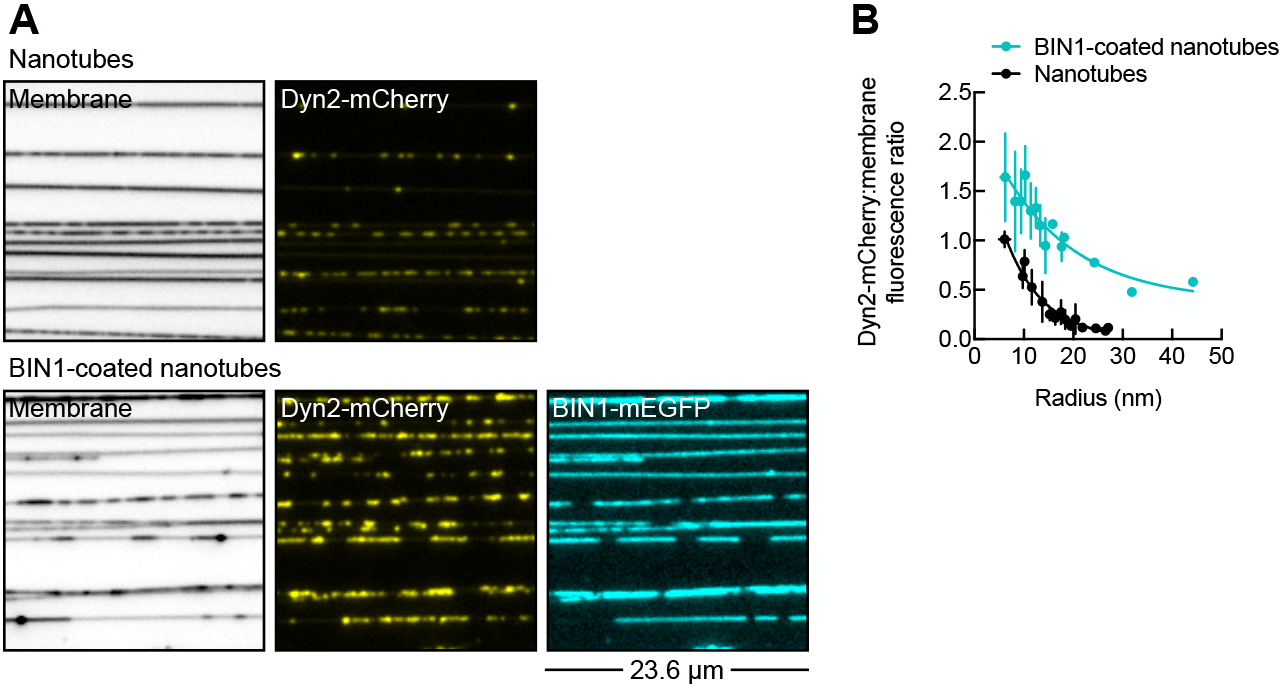
(A) Fluorescence images showing Dyn2-mCherry on nanotubes (top) or BIN1-coated nanotubes (bottom). Nanotubes were exposed to 0.2 μM BIN1-mEGFP and excess protein was washed off. Nanotubes were then incubated with excess 4.6 μM Dyn2-mCherry with 1 mM GppNHp (Jena Bioscience), incubated for 10 mins, washed and imaged. PI(4,5)P_2_ concentration was brought down from 5 to 1 mol% to assess BIN1-dependent Dyn2 recruitment. Membrane is colored in gray and inverted in contrast. (B) Plot showing Dyn2 membrane density measured as the ratio of Dyn2-mCherry and membrane fluorescence on nanotubes of varying sizes with or without BIN1. Each data point represents the mean ± SD of Dyn2 membrane density on tubes binned to their integral radius values.

